# A minimally invasive EEG recording method in mice using thin needle electrodes

**DOI:** 10.64898/2026.03.31.715731

**Authors:** Bende Zou, Xinmin (Simon) Xie, Ludmila Gerashchenko

## Abstract

Currently, implantation of electroencephalogram (EEG) electrodes in laboratory animals is time-consuming and requires specialized equipment. We present a novel method for EEG recordings in mice that utilizes thin needle electrodes. These electrodes are inserted into the skull at predetermined locations by gently pressing them against the bone surface. To ensure stable fixation of the implant, hook-shaped needles are positioned along the lateral aspects of the skull. The electrodes are connected to a multipin connector and secured to the skull using dental composite, after which the animal is allowed to recover from anesthesia. Importantly, procedures such as skull drilling and screw placement are not required, allowing the entire surgery to be completed in less than 15 minutes. Consequently, this EEG implantation approach is rapid and minimally invasive. Results of our studies indicate that EEG recordings obtained with needle electrodes are not inferior to those obtained with screw electrodes. Overall, the method is designed to enhance the accuracy and efficiency of EEG recording studies while improving animal welfare.

Simplifies the placement of EEG electrodes.
Reduces the time required for electrode implantation.

**Graphical abstract:** 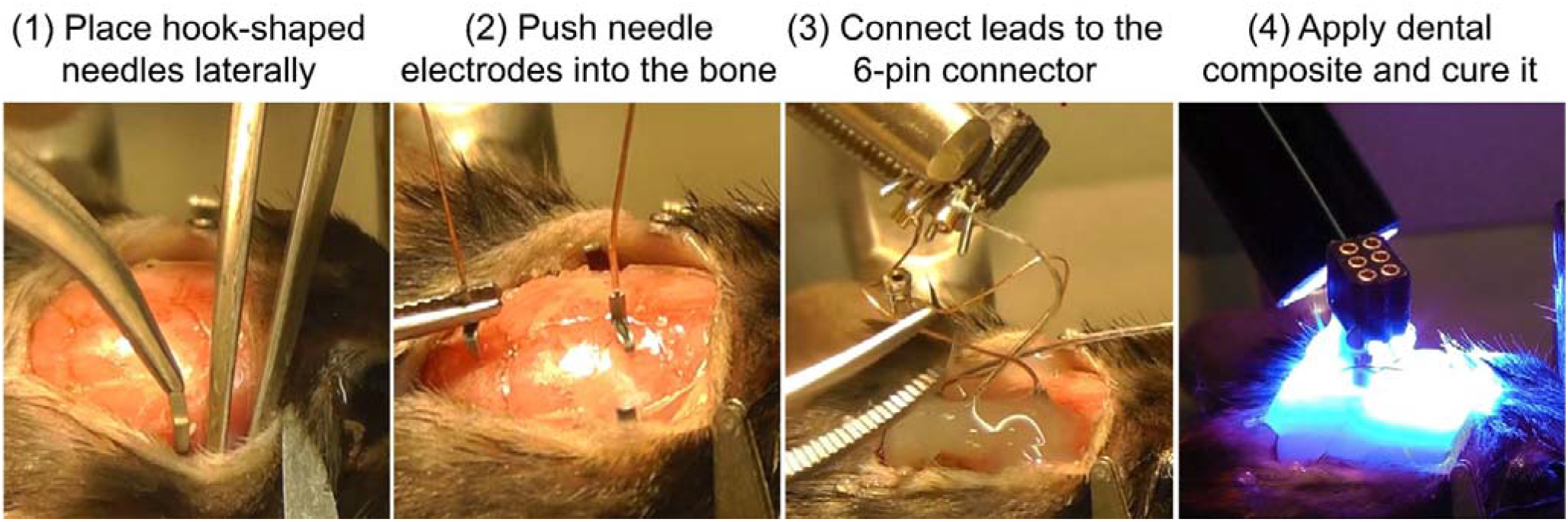

## Background

Electroencephalography is widely used to measure the brain’s electrical activity. This technique is commonly applied in animal studies of epilepsy,[1] sleep–wake cycles,[2] brain injury and neurological disorders,[3] and pharmacological responses.[4]

Currently, implantation of electroencephalogram (EEG) electrodes in laboratory animals is time-consuming and requires specialized equipment. Electrode placement is typically performed in anesthetized animals by drilling a hole in the skull at the desired location using a stereotaxic apparatus. In most cases, screw-type EEG electrodes are used; however, these may produce lesions in the underlying cortex. Tissue damage can occur when screw insertion exceeds skull thickness (the average thickness of the mouse skull is approximately 0.2 mm) and compresses or penetrates the dura mater. Injury to the dura mater or cortex may lead to neuroinflammation and gliosis, resulting in scar tissue formation around the electrode. Such changes can alter electrode impedance, degrade EEG signal quality, and affect subsequent recordings even after prolonged recovery periods. Furthermore, implantation of EEG electrodes in laboratory animals typically requires at least 30–60 minutes, even when performed by an experienced investigator. Prolonged surgical duration—particularly extended anesthesia—places additional physiological stress on the animal and may impair postoperative recovery, potentially increasing morbidity or mortality. In addition, the financial cost of EEG implantation is substantial, including expenses associated with stereotaxic equipment, surgical supplies, dental cement, screw electrodes, inhalational anesthesia, and use of a dedicated surgical facility. Therefore, improving time efficiency, simplifying surgical procedures, and minimizing brain damage associated with screw electrode placement in small laboratory animals such as mice are highly desirable objectives.

Several approaches have been developed to reduce brain injury during EEG recordings, including subdermal needle electrodes[5] and cortical surface electrode arrays.[6,7] Although nonpenetrating methods preserve brain tissue for subsequent histological analysis, they often yield lower signal quality and reduced spatial precision.[8] Conversely, electrodes that penetrate the skull and enter the brain generally provide clearer signals and more accurate localization but are more invasive. The use of ultrathin penetrating electrodes may allow both high signal quality and good tissue preservation. Here, we describe novel tools and procedures for the placement of needle-based EEG electrodes that are rapid and minimally invasive. This approach aims to increase the accuracy and efficiency of EEG studies while improving animal welfare.

## Method details

The components recommended for needle electrode placement are listed in Table 1. While we used these components in our experiments, some may be replaced with equivalent items commonly available in other research laboratories (e.g., acrylic cement may replace flowable composite). Certain procedural steps are optional; our experience indicates that the strongest and most durable fixation is achieved when both Optibond and hook-shaped needles are used. However, sufficient fixation can be obtained with hook-shaped needles alone. Because bone drilling is unnecessary, and ear bar placement may also be omitted, surgical time can be significantly reduced.

**Table 1.**
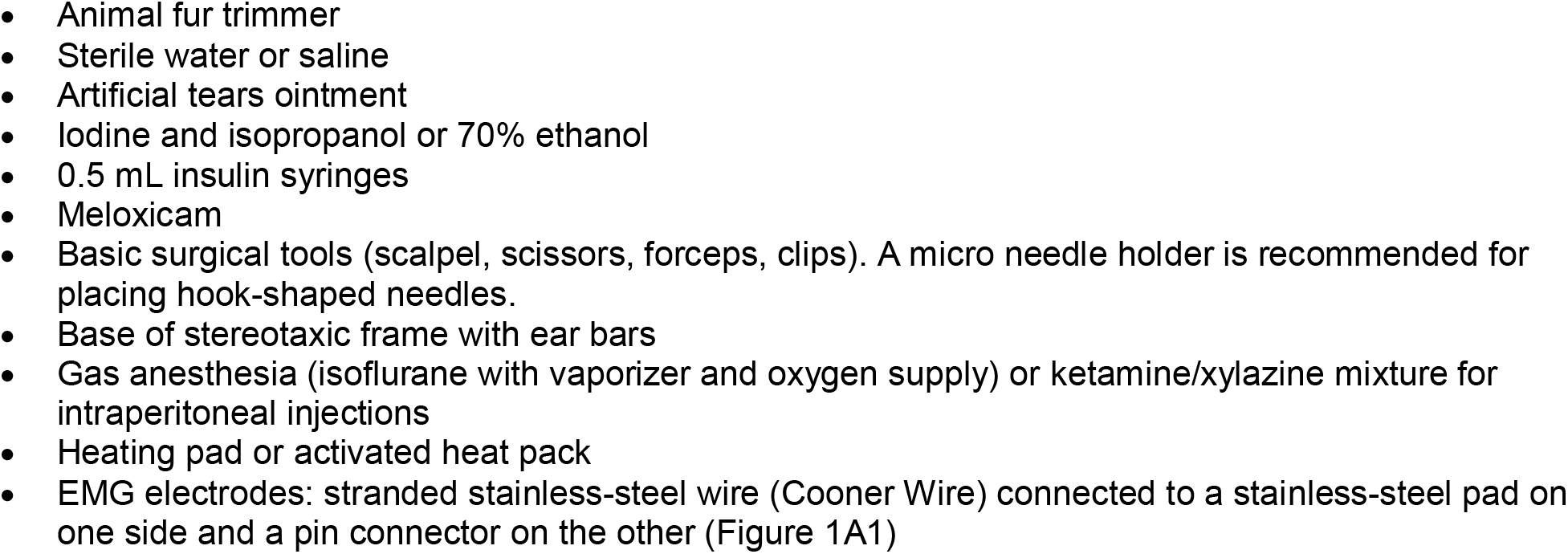
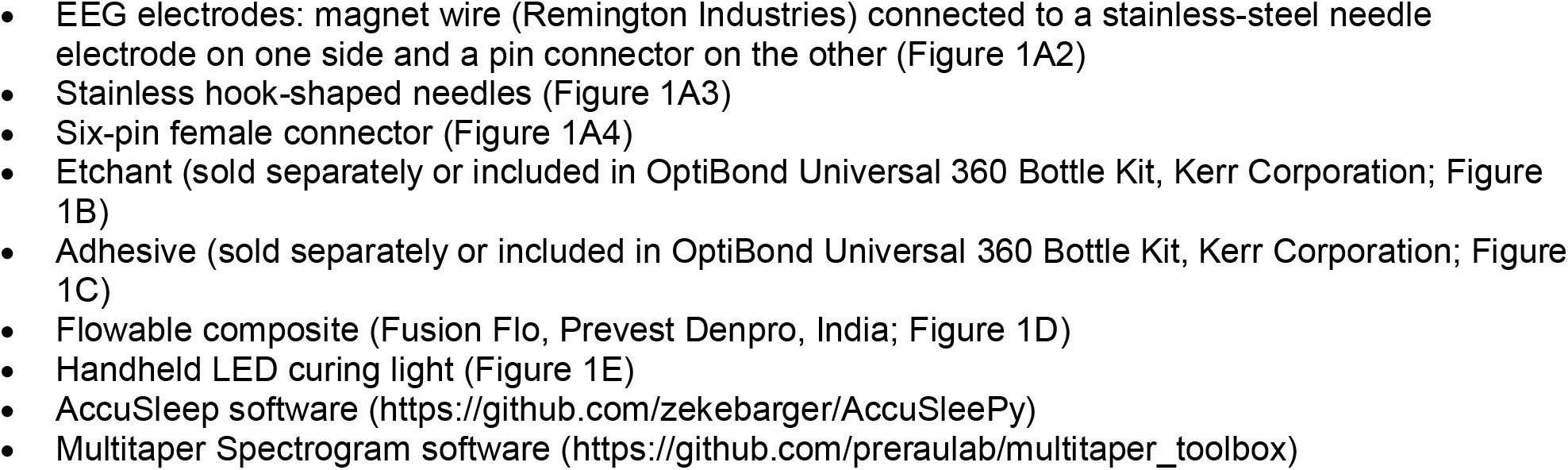
Equipment list.

**Figure 1.**
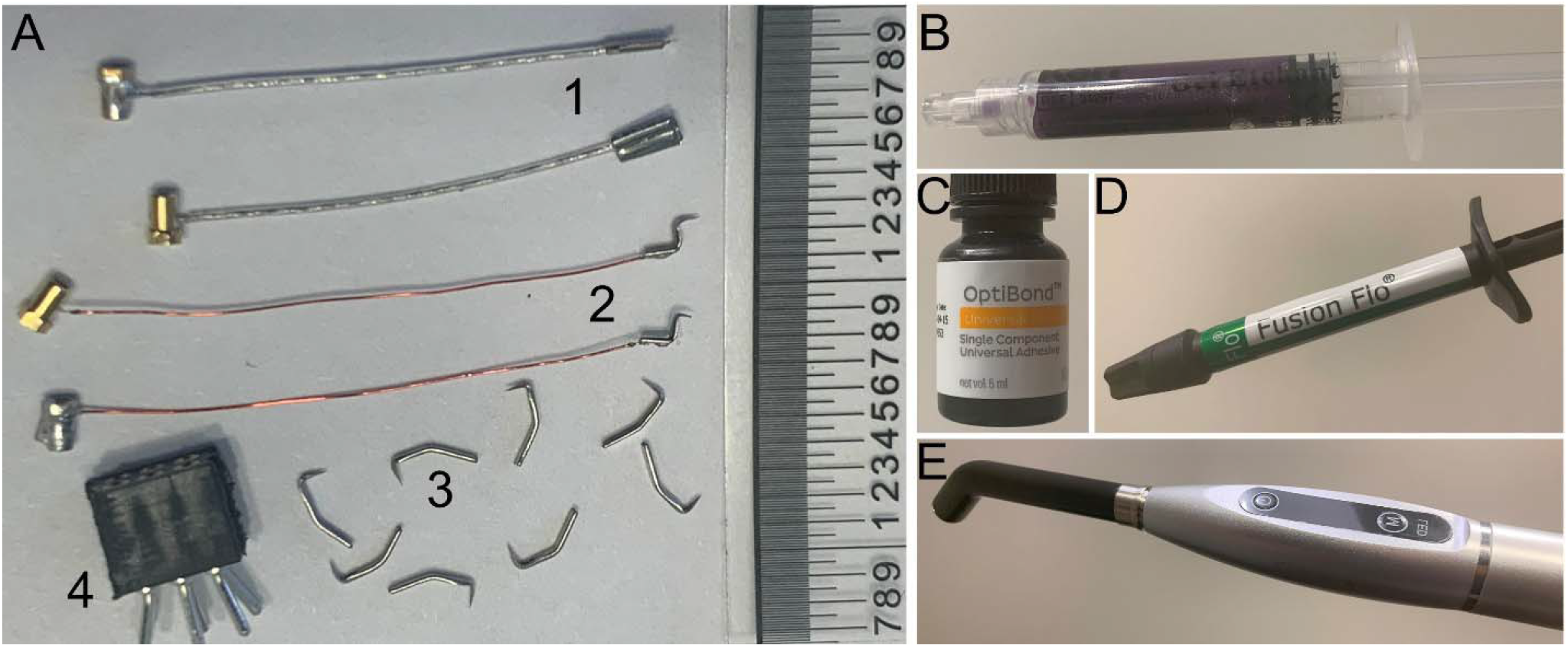
Components required for placement of needle electrodes. EMG electrodes (A1) and needle electrodes (A2) are soldered to female connectors that can be connected to the desired lead of the 6-pin connector (A4). Hook-shaped needles (A3) are used to secure the implant to the skull. The etchant (B), Optibond (C), composite (D), and LED lamps are used during surgery.

The procedure for electrode placement is as follows. These steps are also demonstrated in the accompanying videos:

1. Anesthetize the mouse with 1.5–2% isoflurane in oxygen or by intraperitoneal (i.p.) injection of ketamine (100 mg/kg) and xylazine (10 mg/kg).
2. Fix the mouse’s head in the base of a stereotaxic instrument.
3. Place a heating pad or activated heat pack under the mouse to maintain body temperature.
4. Remove hair from the top of the head using an electric clipper or scissors.
5. Disinfect the scalp by wiping it with iodine, followed by ethanol or isopropanol using a cotton swab.
6. Make a midline skin incision of approximately 1 cm with a scalpel. Use two clips to retract the skin and expose the skull.
7. Scrape soft tissue from the top of the skull using a scalpel.
8. Apply a small drop of an etchant (phosphoric acid) from the Optibond kit onto the skull. Distribute it evenly with the tip of the syringe and leave for 30–60 seconds.
9. Remove the phosphoric acid with a cotton swab and rinse the skull three or four times with sterile water or 0.9% NaCl. **Note:** Phosphoric acid must be completely removed to prevent tissue damage.
10. Dry the skull with a cotton swab and compressed air. **Note:** If compressed air is not available, allow the skull to air-dry for 1 minute.
11. Apply a small drop of Optibond over the skull using a disposable brush provided in the kit. Cure the Optibond for 10 seconds using blue light. **Note:** Ensure the skull is completely dry before applying Optibond. Avoid direct eye exposure to the blue light.
12. The combination of etchant, Optibond, and light-curable composite is typically sufficient to secure the implant for one to two weeks. For stronger fixation or if Optibond is not used, miniature hooks can be placed along the sides of the skull. Detach the muscle from the skull using miniature forceps (Figure 2A). Hold the hook-shaped needle with a micro needle holder and gently insert it into the bone (Figure 2B and 2C). **Note:** In young mice, only light pressure is required because the lateral skull is thin. Excessive force may indicate that the needle holder, rather than the needle, is pressing against the bone. Greater pressure may be required in older mice with thicker skull bones.
13. Position the needle EEG electrode at the desired location and gently push it into the bone (Figure 2D and 2E).
14. Apply fluid dental composite (e.g., Fusion Flo) around the implant and surrounding area. Cure for 10–20 seconds with blue light.
15. Insert EMG electrodes between the skin and neck muscles. Add additional composite around the electrodes and cure with blue light. **Note 1:** EMG electrode placement may be omitted if SmartCage™ (AfaSci, Inc. Burlingame, CA, USA) is used in the study (Figure 3). SmartCage™ is a modular platform that records rodent activity and behavior using sensors placed under or around the cage.[9] **Note 2:** Sufficient composite is essential to firmly secure the implant, as a loose fixation may result in movement artifacts in EEG recordings.
16. Connect EEG and EMG electrodes to the 6-pin connector. Add composite around the connector and cure with blue light.
17. Administer a subcutaneous analgesic (meloxicam, 5 mg/kg) and place the mouse on a heating pad until it recovers from anesthesia and resumes normal movement in the cage.

**Figure 2.**
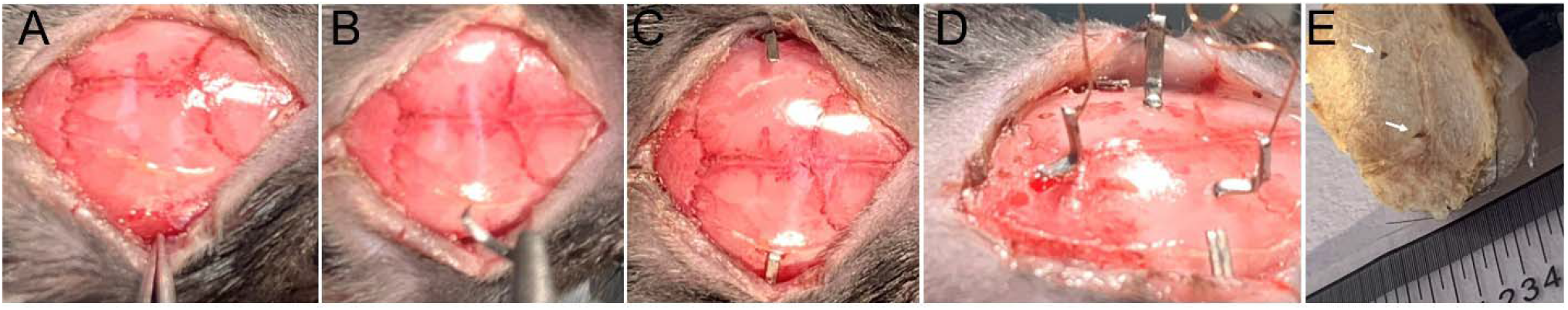
Placement of hook-shaped needles and needle electrodes in the mouse skull. The lateral muscle is displaced ventrally by approximately 1–2 mm (A). The hook-shaped needle is held with a micro needle holder (B) and gently inserted by pressing it into the bone (C). Needle electrodes are positioned at the desired locations and pushed into the bone (D). Ventral view of the removed implant indicates that the needles penetrate the bone into the brain by less than 0.5 mm (E).

**Figure 3.**
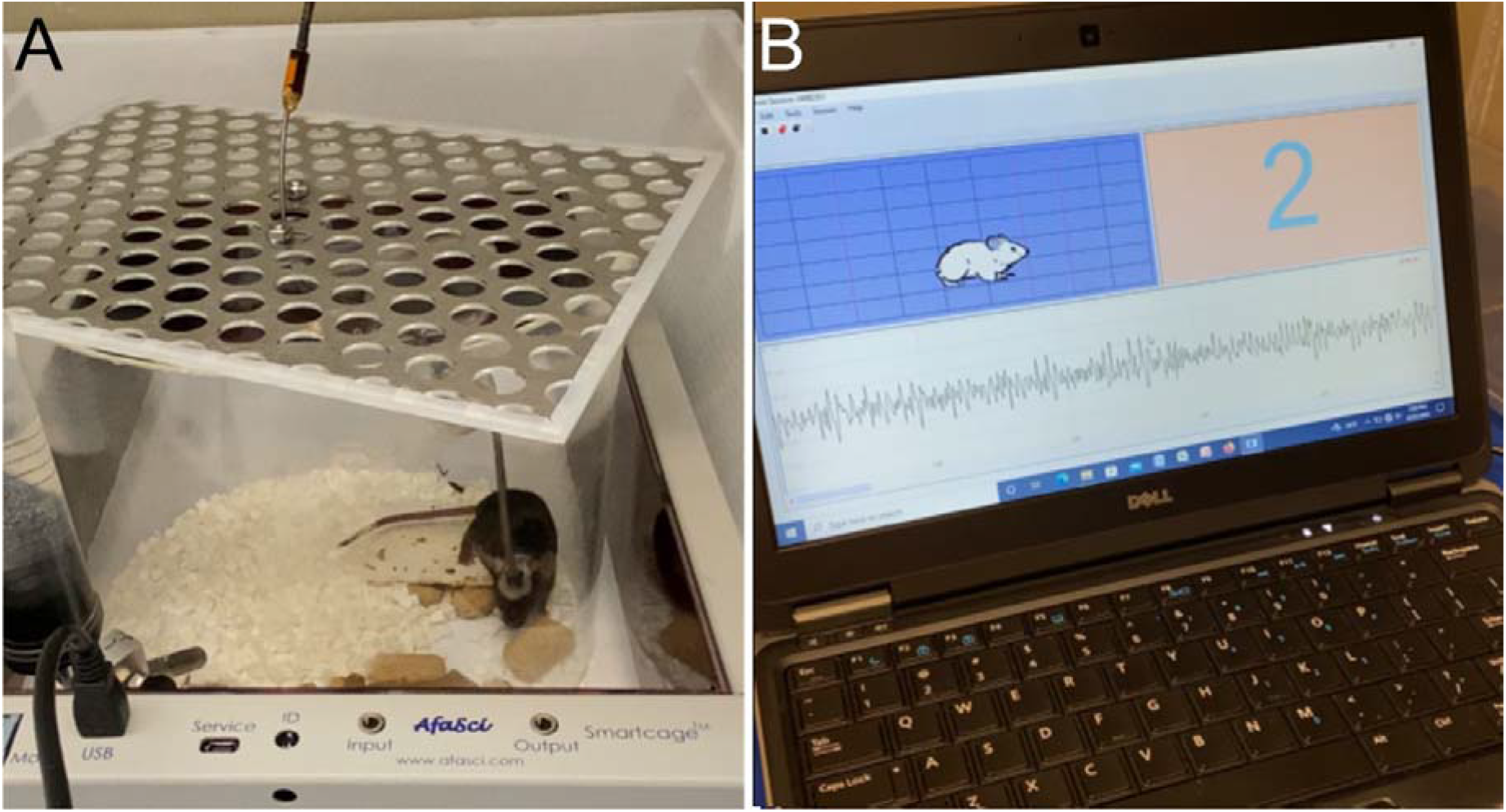
Simultaneous recording EEG and mouse activity using SmartCage™. The SmartCage™ platform provides an inner space of 36.0 × 23.0 × 9.0 cm (length x width x height) for a regular mouse cage (A). Home cage activity variables, including activity counts (counts of breaks in x-, y- and z-axis photo beams), activity time, locomotion (distance travelled and speed) and rearing, are calculated using CageScore™ software (B).

## Method validation

To validate the method, we compared EEG recordings in mice implanted with screw electrodes and needle electrodes.

### Animals

C57BL/6J mice (stock no. 000664) were obtained from The Jackson Laboratory and bred in-house. Mice were housed at 70–74°F under a 12-h light/dark cycle (lights on 7:00 AM–7:00 PM) with food and water available *ad libitum*. At 3–5 months of age, mice were implanted with EEG and EMG electrodes and used for EEG/EMG recording studies.

### Regular implantation procedure: screw electrodes

Six male mice were used for this experiment. Mice were deeply anesthetized with isoflurane (1–3%) and secured in a stereotaxic frame. The scalp was incised and retracted to expose the skull. An etchant was applied to the skull surface for 30–60 seconds, followed by rinsing with sterile water and brief drying. A thin layer of OptiBond was applied to the skull and cured using blue light for 10–20 seconds. An EEG electrode was screwed into the skull above the parietal cortex (anteroposterior = −1.0 mm, mediolateral = 2.0 mm). A reference screw electrode was placed above the cerebellum (1.0 mm posterior to lambda).

Electromyogram (EMG) electrodes were inserted into the nuchal muscles. All electrodes were connected to a 6-pin connector and secured to the skull using a light-curable composite. Mice received meloxicam for postoperative analgesia and were allowed to recover.

### Novel implantation procedure: needle electrodes

Six male mice were anesthetized with 1.5–2% isoflurane in oxygen and placed in a stereotaxic frame. The scalp was shaved and incised to expose the skull. An etchant was applied to the skull surface for 30–60 seconds, followed by rinsing with sterile water and brief drying. A thin layer of OptiBond was applied to the skull and cured using blue light for 10–20 seconds. Two hook-shaped needles were positioned along the lateral aspects of the skull to improve implant stability. Needle EEG electrodes were then gently pressed into the parietal and occipital bone (recording electrode: anteroposterior = −1.0 mm, mediolateral = 2.0 mm; reference electrode: 1.0 mm posterior to lambda). The needles penetrated the skull without the need for excessive force and without causing bleeding. The electrodes were secured using a light-curable composite. EMG electrodes were inserted subcutaneously into the nuchal muscles. All implant components were further stabilized with additional light-curable composite. Mice received meloxicam for postoperative analgesia.

### In vivo EEG/EMG recordings

After at least one week of postoperative recovery and two days of habituation to the recording chamber, mice were tethered to the EEG/EMG recording system and allowed to move freely in their cages. EEG and EMG signals were amplified and filtered using a Grass Neurodata Model 12 Amplifier system (Grass Instrument Division of Astro-Med, Inc., West Warwick, RI). Signals were digitized at 512 Hz. EEG signals were filtered with low- and high-frequency cutoffs of 0.3 and 30.0 Hz, respectively. Data were stored for subsequent analysis.

### Sleep pattern analysis

Data files were imported into AccuSleep software[10] for automated scoring of sleep states in 5-second epochs over a 24-hour recording period. Vigilance states—including wakefulness, non–rapid eye movement (NREM) sleep, and rapid eye movement (REM) sleep—were quantified in 1-hour bins. Wakefulness was characterized by low-amplitude EEG and high, variable EMG activity. REM sleep was identified by low-amplitude EEG dominated by theta (θ) activity (5–10 Hz in rodents) and muscle atonia with occasional twitches. NREM sleep was defined by high-amplitude, slow-wave EEG and low-amplitude EMG activity. The total time spent in wakefulness, NREM sleep, and REM sleep was plotted in 1-hour bins as a function of Zeitgeber time (ZT).

### Power spectral density analysis

EEG signals from each 5-second epoch were subjected to multitaper spectral analysis. Power spectra were calculated over the 24-hour recording period and analyzed separately for wakefulness, NREM sleep, and REM sleep. Spectral analysis was performed using a Python implementation of the multitaper script developed by the Prerau Lab.[11] EEG datasets were processed separately for each vigilance state using the following parameters:

fs = 512 # Sampling Frequency

frequency_range = [0, 30] # Limit frequencies from 0 to 30 Hz

time_bandwidth = 3 # Set time-half bandwidth

num_tapers = 5 # Set number of tapers

window_params = [5, 5] # Window size is 5s with step size of 5s

min_nfft = 0 # No minimum nfft

detrend_opt = ‘constant’ # detrend each window by subtracting the average

multiprocess = True # use multiprocessing

cpus = 3 # use 3 cores in multiprocessing

weighting = ‘unity’ # weight each taper at 1

plot_on = True # plot spectrogram

return_fig = False # do not return plotted spectrogram

clim_scale = True # do not auto-scale colormap

verbose = True # print extra info

xyflip = False # do not transpose spect output matrix

The program generated spectrogram matrices spanning 0–30 Hz with a frequency resolution of 0.125 Hz. Outliers exceeding 10× the standard deviation within each frequency bin were removed prior to further analysis. To account for interindividual variability in absolute EEG power, power spectral density (PSD) values for each frequency bin and vigilance state were normalized to a reference value calculated for each mouse. The reference value was defined as the mean total EEG power across all frequency bins (0.5–30 Hz) and all behavioral states over the 24-hour period. Normalized EEG power was expressed as a percentage of total power within the 0.5–30 Hz range and presented in 0.125-Hz bins across the 0.5–30 Hz frequency range for the 24-hour recording period.

### Statistical analysis

In all behavior tests and scoring, the researchers were blinded to treatments. All data are presented as mean ± standard error of the mean (SEM). Because our experiments involved two groups with two factors (treatment and time), two-way analysis of variance (ANOVA) for repeated measures was used for overall comparisons. If significant differences were found for the main effects or interactions, post hoc analyses using Fisher’s protected least significant difference (PLSD) test, or Tukey test, were conducted.

## Results

Although the surface area occupied by screw electrodes is substantially larger than that of needle electrodes, the EEG signals (Figure 4A–B) and power spectral density (Figure 5C–D) were nearly identical between the two electrode types. ANOVA did not show significant differences between groups at any timepoint of the study. A likely explanation of this result is that EEG signal power depends primarily on the relative locations of the recording and reference electrodes, which were similar for both screw and needle configurations. Furthermore, no significant differences were observed in the amounts of wakefulness, REM sleep, or NREM sleep between the two recording methods (Figure 5A–B). These results indicate that EEG recordings obtained with needle electrodes are not inferior to those obtained with screw electrodes.

**Figure 4.**
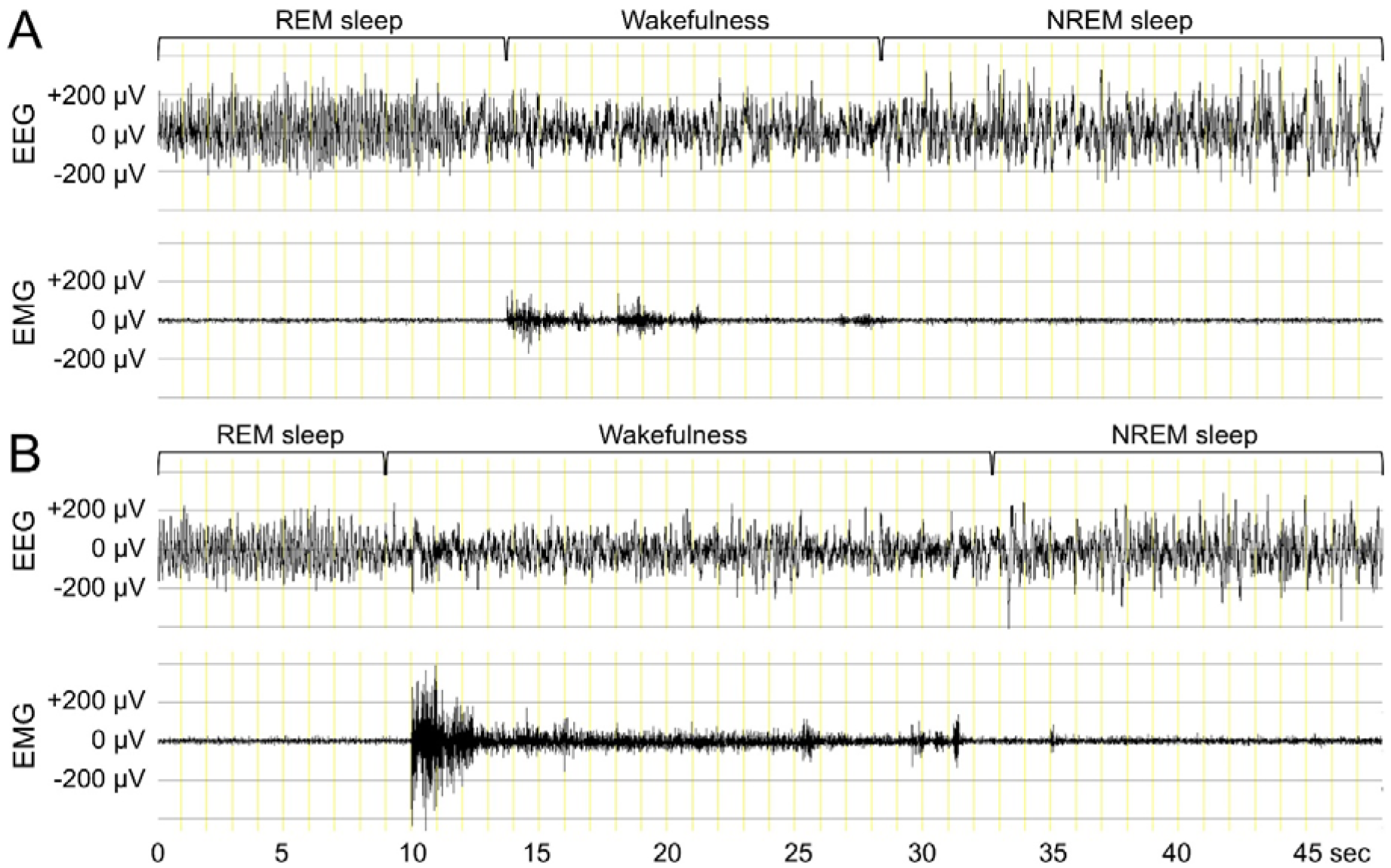
Representative EEG and EMG recorded in a mouse using screw electrodes (A) or needle electrodes (B) during transitions from REM sleep to wakefulness to NREM sleep.

**Figure 5.**
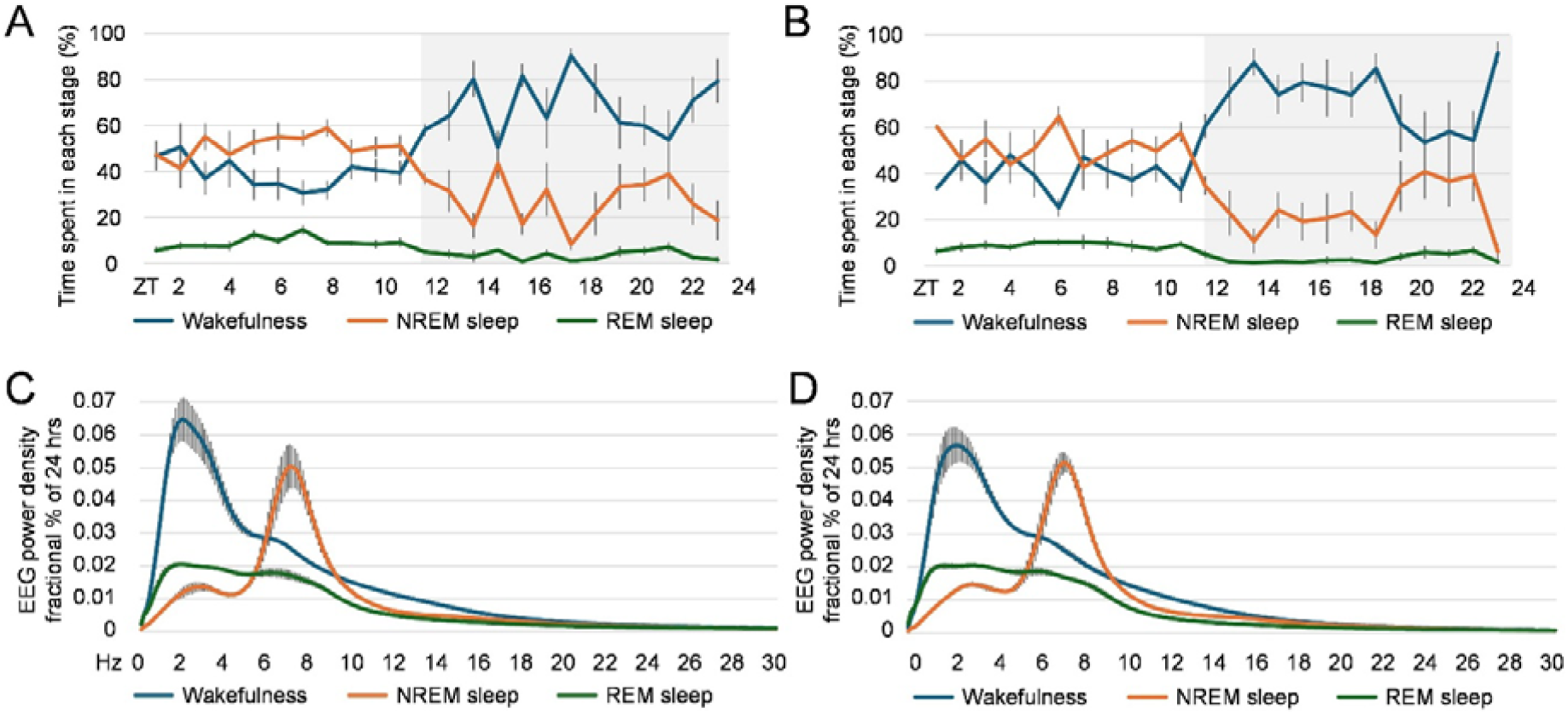
Amounts of wakefulness, REM sleep, and NREM sleep (A–B) and power spectral density (C–D) recorded in mice using either screw electrodes (A, C) or needle electrodes (B, D) over a 24-hour period. Shaded areas in A and B indicate the light-off period. Six mice were included in each electrode group. Data are presented as mean ± SEM.

## Limitations

Screw electrodes are commonly used despite their potential to cause brain damage and, in some cases, animal loss. Published studies report mortality rates of 15% or higher, increased numbers of epileptiform discharges, alterations in the brain fluid transport system, and direct brain injury in mice implanted with screw electrodes.[12-15] Approximately 12–24 hours after screw implantation surgery, mice often stop moving and eating and become hypothermic. These animals frequently require prolonged supportive care, including heating pads, palatable food, and intraperitoneal saline injections to prevent dehydration. Despite these interventions, some mice do not survive screw implantation procedures. Hemorrhage—either subdural or intraparenchymal—resulting from screws penetrating the brain tissue is a likely cause of these adverse outcomes.

In contrast, none of these complications were observed following needle electrode implantation. Animals recovered rapidly and displayed normal home-cage behavior, eliminating the need for heating pads or specialized post-operative care. In our experiments, mice implanted with needle electrodes resumed eating, drinking, and exhibiting normal behavior immediately after recovery from anesthesia.

Besides screw electrodes, anchor screws are widely used in mouse neurosurgery to provide stable mechanical fixation for cranial implants, including EEG/EMG electrodes, headstages, optical fibers, and cannulas. These screws serve as anchoring points for dental cement or composite materials, reducing the risk of implant detachment during chronic recordings and enabling experiments in freely moving animals. However, screw placement increases surgical time and the risk of brain injury and inflammation, particularly in young or fragile mice. We have developed hook-shaped needles specifically for use in mice and other small laboratory animals to eliminate the need for anchor screws. Hook-shaped needles provide stronger fixation of implants to the skull than other existing methods. This stability is achieved by positioning the hooks along the lateral aspects of the skull, ensuring that the implant remains secure unless the metal hooks themselves fail. Because screws can cause significant brain damage, replacing them with anchor needle hooks represents a desirable alternative for research applications.

In conclusion, needle EEG electrodes offer several advantages for neuroscience research:

1. Improved animal welfare: Needle electrodes do not cause significant brain damage, and implantation has not resulted in mortality.
2. Improved data quality: Brain damage caused by screw electrodes may confound experimental results; this issue is avoided with needle electrodes.
3. Simplified and faster surgical procedures: Bone drilling and screw placement are unnecessary, reducing surgical time and labor. Needle electrodes can be implanted in less than 15 minutes, whereas screw electrode implantation is more complex and typically requires 30 minutes or longer, even for experienced surgeons.

Several limitations of using needle EEG electrodes should be mentioned. Because needle EEG electrodes sit in the skull, they record spatially broad field potentials from the cortex rather than activity from precise locations, and signals from deep brain structures are largely attenuated. In addition, only a small number of needle electrodes can be placed on the mouse skull, which limits spatial sampling of cortical activity, and recordings can be affected by reference contamination, muscle activity, and movement artifacts.

Consequently, needle EEG electrodes are best suited for monitoring global brain states rather than detailed spatial or circuit-level neural activity.

## Supporting information

Video 1- Placement of hook-shaped needles and needle electrodes

Video 2 - Complete procedure

## Ethics statements

All procedures were conducted in accordance with the National Institutes of Health guidelines and were approved by the AfaSci Institutional Animal Care and Use Committee. The study adhered to the updated Animal Research: Reporting of In Vivo Experiments (ARRIVE 2.0) guidelines.

## CRediT author statement

Bende Zou: Methodology, Investigation, Formal analysis, Writing – review & editing. Xinmin (Simon) Xie: Conceptualization, Methodology, Resources, Supervision, Writing – review & editing. Ludmila Gerashchenko: Conceptualization, Methodology, Resources, Investigation, Formal analysis, Supervision, Funding acquisition, Writing – original draft, Writing – review & editing.

## Acknowledgments

This work was supported by the National Institutes of Health (NIH) grant R43MH135817 (former number R43OD035215).

## Declaration of interests

The needle electrodes and hook-shaped needles used in this study were developed by Neurotargeting Systems, Inc. Ludmila Gerashchenko, the founder and owner of Neurotargeting Systems, holds the patent for the production and use of these needle electrodes and hook-shaped needles. This invention was made with government support under the NIH grant, and the government has certain rights in the invention. Xinmin (Simon) Xie, the founder and owner of AfaSci, Inc., holds the rights for SmartCage™ and CageScore™.

## Supplementary material

Supplementary videos associated with this article can be found in the online version.

